# Memory shapes microbial populations

**DOI:** 10.1101/2020.11.05.370106

**Authors:** Chaitanya S. Gokhale, Stefano Giaimo, Philippe Remigi

## Abstract

Correct decision making is fundamental for all living organisms to thrive under environmental changes. The patterns of environmental variation and the quality of available information define the most favourable strategy among multiple options, from randomly adopting a phenotypic state to sensing and reacting to environmental cues. Memory – a phenomenon often associated with, but not restricted to, higher multicellular organisms – can help when temporal correlations exist. How does memory manifest itself in unicellular organisms? Through a combination of deterministic modelling and stochastic simulations, we describe the population-wide fitness consequences of phenotypic memory in microbial populations. Moving beyond binary switching models, our work highlights the need to consider a broader range of switching behaviours when describing microbial adaptive strategies. We show that multiple cellular states capture the empirical observations of lag time distributions, overshoots, and ultimately the phenomenon of phenotypic heterogeneity. We emphasise the implications of our work in understanding antibiotic tolerance, and, in general, survival under fluctuating environments.

## Introduction

In an ideal world, living organisms would be able to adapt instantly and reliably to changing environmental conditions in order to maximise their instantaneous performance. However, conditions may change abruptly and unpredictably, making it ineffective to mount a specific rapid response. Also, some responses require the synthesis of complex molecules (secretion systems or capsules in bacteria) or entering a physiological state (dormancy) that cannot be reverted instantaneously, if need be. Switching to a new phenotype may thus commit the cell to a response that lasts for a specific timescale different than the duration of the changed environment. Besides phenotypic switching, diversity in response rates can result in intricate patterns of phenotypic heterogeneity. We postulate that instantiating memory in switching mechanisms underpinning phenotypic heterogeneity will affect population composition, size and ultimately fitness. While classical models of phenotypic heterogeneity use simple on-off dynamics, recent results stemming from accurate, modern experimental methods require us to delve deeper into the dynamics. Here we provide a theoretical framework for phenotypic switching rooted in a mechanistic concept of molecular memory.

In bacteria, memory may emerge as a component of phenotypic heterogeneity [1,2,3,4,5]. In homogeneous environments, non-genetic individuality can arise through fluctuations in molecular concentrations of signaling molecules during transcriptional bursts [6], unequal partitioning at cell division [7] or via other epigenetic mechanisms [8,9,10, 11, 12]. Such ‘in-built’ mechanisms for phenotypic variation help bacterial populations adapt to harsh environments [13, 14, 12, 15]. When environmental fluctuations show stereotypical patterns, unicellular organisms may harness this temporal information to adjust their mode of phenotypic adaptation to match, or even anticipate, environmental fluctuations [16]. Such strategies can be embedded in genetic regulatory networks [17, 18] or arise from epigenetic phenotypic switches [19, 20]. These forms of fitness optimization by associative learning can arguably be assimilated to memory-based processes [21, 22].

A large body of theoretical work recognises the role of phenotypic heterogeneity by analysing its effect on bacterial fitness [14, 23, 24, 25]. However, depending on the details of the underlying genetic network controlling switching, its dynamics may not be adequately described by simple ‘on’ -‘off’ dynamics with constant rates of switching between distinct phenotypic states. When phenotypic switching occurs at a constant rate, the residence time in a given phenotypic state is best described mathematically as a memory-less process [26]. The probability of switching, then, only depends on current cell state and not on any previous state, leading to exponentially-distributed residence times. By contrast, when the probability of switching depends on time spent in a given state, residence time no longer follows an exponential distribution and multigenerational memory occurs [27]. A simple constant phenotypic switch is, therefore, not a generic model when approaching the problem of intergenerational memory. For example, such a model is inadequate when explaining phenomena such as broad time-lag distributions and both over-and undershooting behaviour of specific phenotypes observed in bacteria or eukaryotic organisms [19, 28, 29].

Motivated by the discovery of different forms of memory in bacteria [27, 26], we propose a unifying approach to define memory. Within the framework of our approach, we ground our model in mechanistic processes occurring in a cell. These processes can be experimentally detected, helping us classify exact phenotypes. First, we derive deterministic dynamics from first principles at the level of individual cells and track the behaviour of a cell population emerging from different switching genotypes. Using this model, we are consistently able to explain the experimental observation of over/undershoots and wide time lag distributions. Then, paying particular attention to the characterisation of transient and equilibrium states of the system, we discuss their consequences on the fitness of a lineage in the presence of fluctuating environments. Significantly, fluctuations across stressed and relaxed environments, as in certain antibiotic treatment regimes, highlight the applicability of our approach. The classically studied bi-stable switch is a special case of our more general model.

## Model & Results

### 2.1 Building memories

We simulate the ecological dynamics of isogenic populations of unicellular organisms that can exist in two different phenotypic states: ‘on’ and ‘off’. Switching from ‘off’ to ‘on’ is unidirectional and stochastic, and occurs at rate *µ*. After turning ‘on’, cells cascade via a deterministic, multi-step process through *n* compartments eventually returning to the ‘off’ phenotype. These compartments represent internal molecular states (potential), such as the dilution of cytoplasmic or membrane protein concentrations [2, 7, 4, 30], that may determine the qualitatively different ‘on’ -‘off’ phenotypes through a threshold-based mechanism [31].

Immediately after turning ‘on’, cells have the highest potential. While retaining the ‘on’ state, cells gradually lose potential by transitioning through the successive compartments at a leaching rate of *ϵ*. This movement reflects the process of protein degradation in a simplistic manner, whereas more complicated forms can be formulated (with bumps or plateaus on the landscape, see SI). While the compartments (*i*) are a representation of the phenotypic dynamics, the growth (birth and death) dynamics of the cells occur separately at rates *b*_*i*_ and *d*_*i*_, respectively.

Fig. 1 visualizes this compartmental model for *n* = 4. The reactions in which an individual cell *X*_*i*_ in compartment *i* is involved are then,

**Figure 1:**
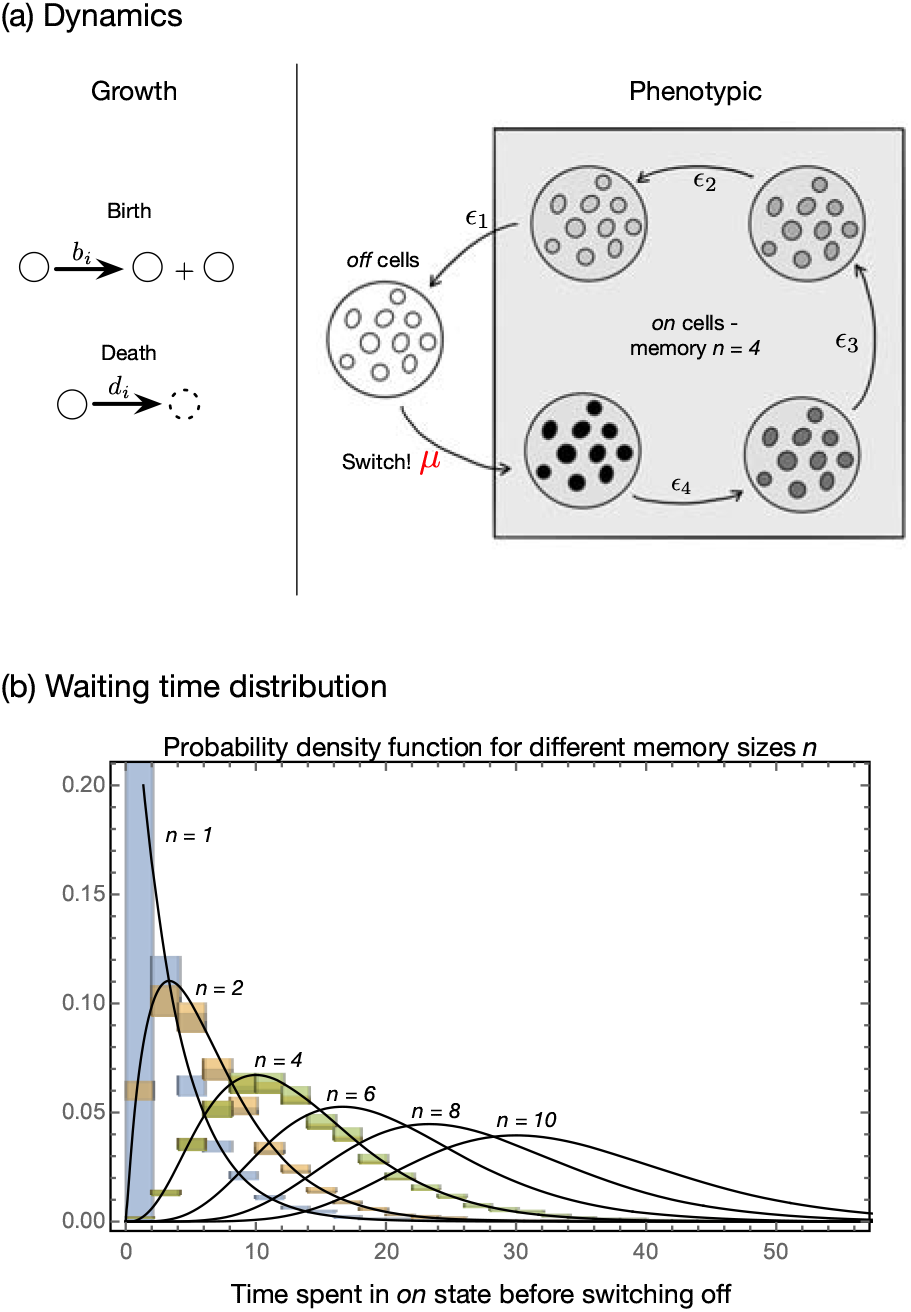
Growth and phenotypic dynamics of ‘off’ and ‘on’ cells. (a) The growth dynamics of cells is given by a birth process and a death process. A cell divides into two cells at rate *b*_*i*_ and dies at rate *d*_*i*_. The index *i* refers to the phenotypic state of being ‘off’ or ‘on’. The ‘off’ cells are the cells in the resting state which is the zeroth compartment *i* = 0. Due to a trigger (an internal constant or a dependence on frequency/density/environmental state) the cells can be turned ‘on’ and they jump to the *n*^th^ compartment (here *n* = 4). The cells do not stay in the ‘on’ state but slowly decay back to the resting state at rate *ϵ* through the different compartments, until the resting state via a number of intermediate phenotypic stages which are different levels of the ‘on’ state. The number of compartments in the ‘on’ state constitutes memory. Thus we intend to make use of this mechanistic interpretation of memory instead of an assumption of a separation between the amount of time a cell spends in ‘on’ or ‘off’ state. (b) Distribution of times a cell stays ‘on’ before switching ‘off’. Starting with a single cell in the completely ‘on’ state we ask how long it takes for the cell to reach the ‘off’ state. Assuming extremely low replication and death rates *b*_*i*_ = *d*_*i*_ = 10^*−*6^, the process is governed by the leaching rates *ϵ* = 0.3. For 1, 2 and 4 ‘on’ compartments we start the Gillespie simulation with one cell in the first ‘on’ state. Once the cells reaches the ‘off’ state we stop the simulation. The normalised histogram of waiting times is then plotted. The probability density function of a gamma distribution with the shape parameter given by the number of ‘on’ compartments and the rate *ϵ* = 0.3 provides the theoretical estimates (black lines) for the simulated 1, 2 and 4 ‘on’ compartments. Increasing the memory, (*n* = 6, 8 and 10) flattens the distribution of the time spent in the ‘on’ state.

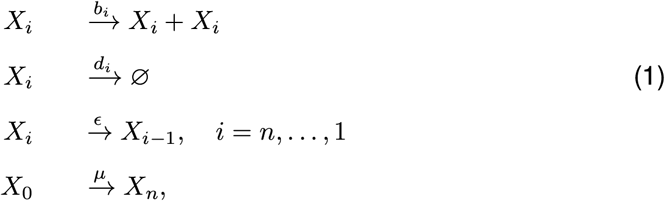

These reactions have a deterministic counterpart in the following linear system of differential equations,

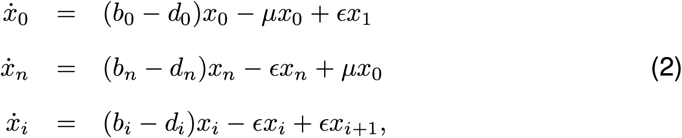

which describe the time evolution of the abundances *x*_*j*_ of cells in the compartments with *i* = 1, …, *n −* 1,. In this model, we can distinguish growth dynamics as specified by birth and death rates from phenotypic dynamics as specified by switching and leaching rates. Their current state determines the movement of cells. Once a cell is in an ‘off’ state it stays ‘off’ until an (external or internal) perturbation switches it back ‘on’. We can imagine the perturbations to be abiotic or biotic [13, 32]. The perturbations have to be large enough to shock the system out of the ‘off’ equilibrium.

In our system, we track the amount of time a cell spends in a particular compartment. Following the trajectories of individual cells, qualitatively different distributions of the time spent in the ‘on’ state emerge as the number of compartments *n* increases. In particular, departure from an exponential distribution is a hallmark of memory [27] (Fig. 1B). For a constant leaching rate, and negligible growth parameters, the length of memory acts like a timer (a deterministic time as termed in [27] for the residence time in the motile state). The overall density function can be captured by a combination of multiple exponential waiting times which results in a gamma distribution with a shape parameter given by the memory size *n* (Fig. 1B). We can thus use this model to study ‘long-term’ memory. Codes for implementing our algorithm as well as for reproducing the relevant figures are available at GitHub.

### 2.2 Asymptotic properties

The asymptotic properties of the model were already well described and do not represent the primary focus of our study [27]. But to make further progress we recapitulate them briefly. The model is Markovian, as the current system state entirely determines cell dynamics. Eventually, as *t* gets very large, the cell distribution in the compartments gets to an equilibrium. This equilibrium can be recovered from an eigenvector of the matrix that captures the system in Eq. 2 (SI). Under complete symmetry in the growth dynamics (*b*_*i*_ = *b* and *d*_*i*_ = *d*) we can get a simple, closed-form expression for this equilibrium for any number *n* of compartments (SI). We begin with a focus on this symmetric case, where the asymptotic growth rate *b − d* of the population is independent of the number of compartments and corresponds to the dominant eigenvalue of the matrix model. The assumption is relaxed further in the manuscript. Our interest is in the effect of changes in the leaching rate *ϵ* and memory *n*. We observe that as *ϵ* decreases or *n* increases, both the time a cell spends in the ‘on’ state and the equilibrium fraction of ‘on’ cells increases (Appendix Fig. SI.1). However, for a small number of compartments, or faster transition through ‘on’ states, the system is dominated by ‘off’ cells.

### 2.3 Transient properties

Indeed, bacteria in natural environments are frequently exposed to changing conditions. Evolution as well as ecological fluctuations can often keep populations from reaching steady states. Thus, a focus on the transient dynamics of eco-evolutionary trajectories is crucial, albeit implying less stringent analytical treatment and greater reliance on computational exploration. While, under constant switching (*µ*), a decrease in *ϵ* is qualitatively equivalent to an increase in *n* in terms of asymptotic properties (i.e. the equilibrium cell distribution and the time spent in either state), this equivalence breaks down when considering transient dynamics. As the number of compartments increases, the frequencies of the ‘on’ and ‘off’ cells do not approach their equilibrium values directly. Instead, the frequencies overshoot, in the case of ‘on’ cells, and undershoot, in the case of ‘off’ cells. The frequency dynamics oscillates when approaching the equilibrium (Fig. 2 a). Interestingly, overshooting and oscillations in the transients of cell frequencies as seen in our model recapitulate observations in cancer cells dynamics [28] and in bacterial growth rate recovery following antibiotic treatments [33]. This transient effect directly relates to the length of memory. While modelling the ‘off’ and ‘on’ states as two compartments would appear more parsimonious, it falls short of replicating distinctive empirical results. Adding compartments intensifies transients, an effect attributable to the spectrum of the matrix model underlying Eq. 2 (see SI). The presence of multiple compartments introduces oscillations due to complex subdominant eigenvalues, absent when there is a single ‘on’ compartment. Increasing the number of compartments also magnifies the influence of complex subdominant eigenvalues on cell dynamics, as the real parts of such eigen-values get closer to the real part to the dominant eigenvalue.

**Figure 2:**
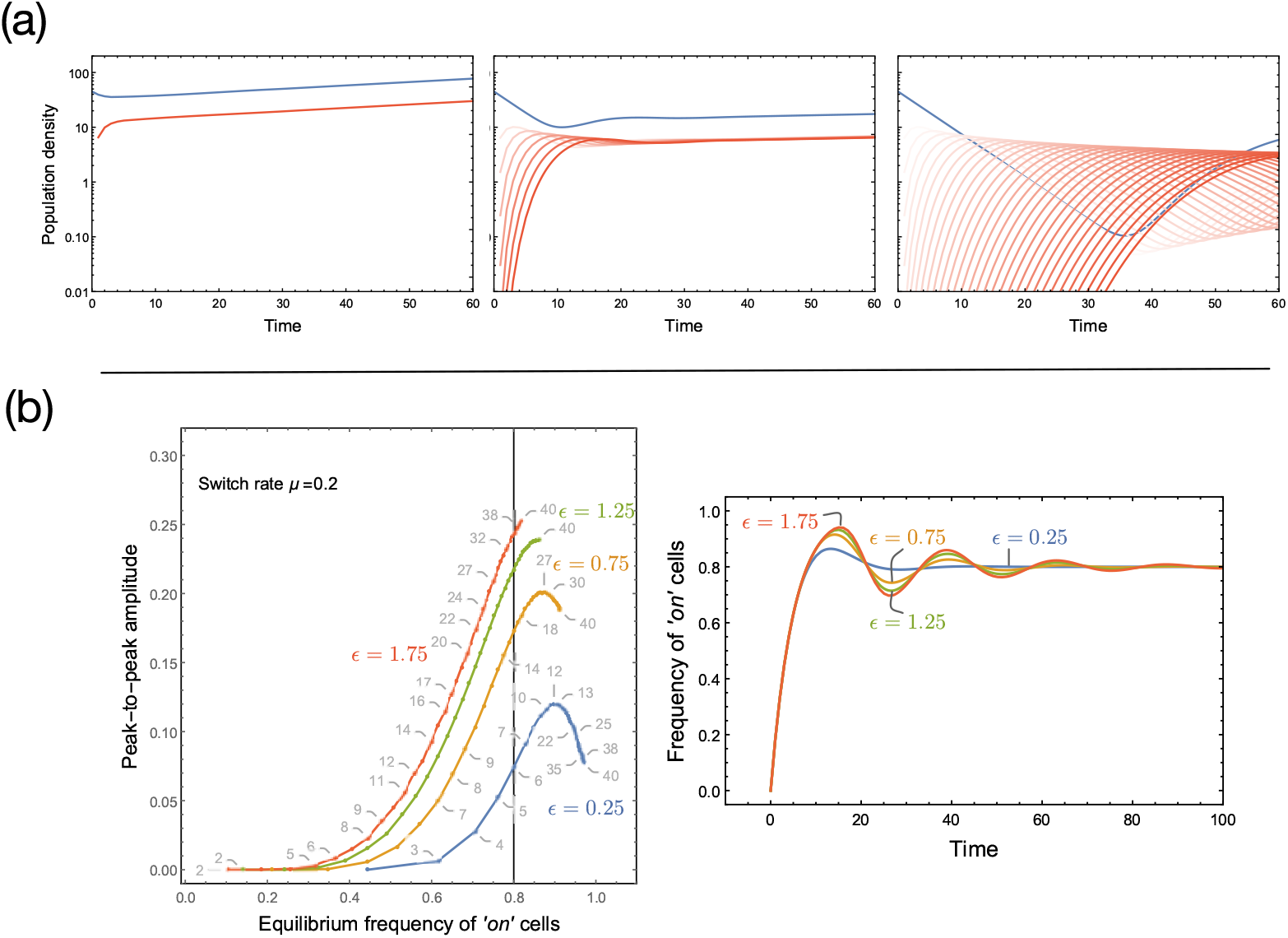
Transient dynamics of multi-state memory. *(a). Multi-state memory and overshoots*. Assuming full symmetry in growth dynamics between the compartments (*b*_*i*_ = *b* = 1.0 and *d*_*i*_ = *d* = 0.98) and a switch rate of *µ* = 0.2 and *ϵ* = 0.5 we show the population dynamics as well as the population composition for fixed time period. The instantaneous growth rate of the two population with different memory sizes is the same *g* = 1*/t*_*max*_ log(*N*_*final*_*/N*_*initial*_). However, with a larger memory, the fraction of ‘on’ cells in the final population is higher. On the way to the equilibrium, multiple compartments result in overshoot dynamics as the ‘on’ cells first need to seed them while depleting the ‘off’ state. *(b). Transient magnitude and number of compartments*. To characterise the oscillations, we plot the peak-to-peak amplitude (the amplitude between the first peak (overshoot) and the second peak (undershoot)) against the equilibrium value of ‘on’ cells as the number of compartments increases (memory (*n*) + 1 (‘off’ cells), gray numbers). The equilibrium fraction of cells in the ‘on’ state is given by *nµ/*(*ϵ* + *nµ*) (SI). Keeping *µ* fixed, this is obtained by varying the leaching rate *ϵ* and by varying the number *n* of compartments in the multi-compartment system, which has constant *ϵ*. As the equilibrium fraction of ‘on’ cells approaches unity, boundary effects prevail, i.e. transient frequency cannot exceed 1, and oscillations’ amplitude gets compressed. The right panel shows the temporal dynamics of ‘on’ cells, all leading to the same equilibrium of 0.8 but for different leaching rate *ϵ* (vertical line in the left panel). The corresponding memory sizes are *n* = 5, 15, 25, 35 respectively. Increasing the number of compartments leads to more pronounced oscillations, not observed in the system with a single compartment regardless of the value of *ϵ* and the equilibrium value of ‘on’ cells. Codes for generating these panels and the general algorithm are available on GitHub.

Cells switch back to the ‘off’ state after spending a certain amount of time in the ‘on’ state. The presence of multiple compartments, extends memory, and affects population composition as well as population density when evaluated at a specific time-point (Fig. 2 a). To capture the transient dynamics when increasing memory, we estimate the peak-to-peak amplitude (the difference between the largest overshoot and undershoot in the frequency of ‘on’ cells) (Fig. 2B). Typically we see a monotonic increase in the relationship between the amplitude and memory length. The relationship wanes as the equilibrium value of ‘on’ cells approaches 1 due to boundary effects Fig. 2B). More generally, the magnitude of the over/undershoot depends on initial conditions and equilibrium of the system.

### 2.4 Environmental variation

While cell lineages can stochastically switch phenotypes even in static conditions [34], such behavior might be evolutionarily advantageous under fluctuating environments [35]. As the environment changes, bacteria can hedge their bets via the well-documented phenomena of persistence [36, 37]. A salient case is bacterial response to different antibiotic treatment strategies.

To explore the effect of memory on bacterial fitness, we consider the case of persisters when subjected to transient antibiotic exposure. Persistence is a common phenomenon in bacteria where a subpopulation of bacterial cells do not grow but is able to survive antibiotic treatments. Persisters can arise stochastically via epigenetic switching, but environmental conditions (nutrient limitation, high cell densities or antibiotic treatments) can also induce their formation (‘triggered’ persistence or type I persisters;[32]. We track population composition and fitness (as proxied by population size) of bacterial lineages under fluctuating environments, consisting of transient exposure to antibiotics. We assign ‘off’ cells to the ‘normal’ physiological state (i.e. growing in permissive conditions, dying under antibiotic treatment). On the other hand, ‘on’ cells are persisters (no growth in either environment and tolerant to antibiotic treatment; see description of parameters in Fig. 3 and example dynamics in the Fig.SI.4). While leaching rate is kept constant, switching from ‘off’ to ‘on’ is triggered only under stress. We consider an initial population of ‘off’ cells, as it represents a stable equilibrium state under conducive growth conditions.

**Figure 3:**
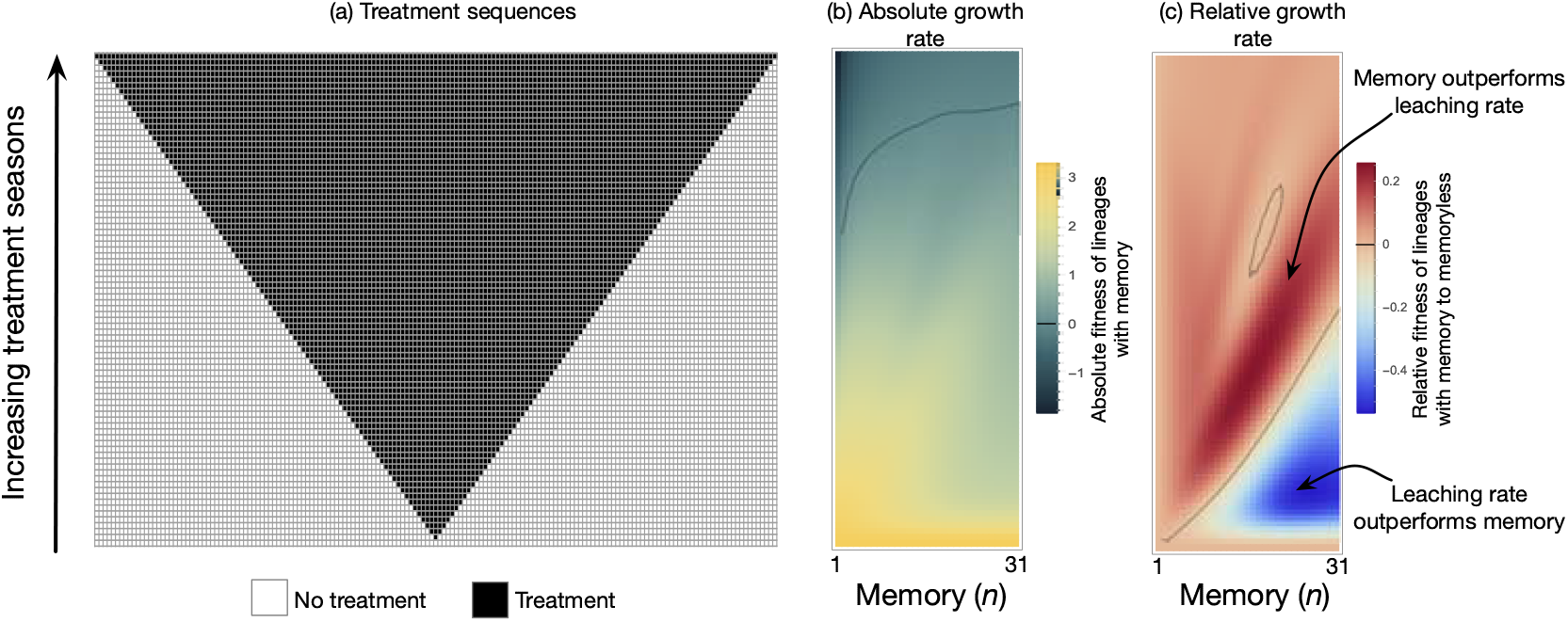
Growth performance of lineages with different memory lengths exposed to treatment of varying length. Cell lineages with different memory sizes (*n* = (1, …, 31) + 1) are temporarily exposed to antibiotic treatment. Each row in the grid is a sequence of seasons from left to right. Seasons with no-treatment (white) and treatment (black) where each lasts for one unit of time. All sequences begin at *t* = 0 with 1000 ‘off’ cells and last until *t*_*max*_ = 165. Under the no-treatment season the growth rate of ‘off’ cells is *b*_*off*_ *− d*_*off*_ = 1 *−* 0.98 = 0.02 and the switch is inactive *µ*_no treatment_ = 0. As cells encounter the treatment season (black squares), the switch is triggered, *µ*_no treatment_ = 0.2 and the death rate of ‘off’ cells increases, *b*_*off*_ *− d*_*off*_ = 1 *−* 1.02 = *−*0.02. The ‘on’ cells are produced but they do not grow, *b*_*off*_ *−d*_*on*_ = 0. The ‘on’ cells leach through the memory compartments at rate *ϵ* = 0.25. At the end of the sequence (*t* = 165), the absolute growth rate, *g*_*m*_ = log (*N*_*memory*_ (*t*_*max*_)*/N*_*memory*_ (0)), was computed. A memoryless process (with an appropriate *ϵ*) can generate the same stable fraction of ‘on’ cells under sustained treatment. We subtract the absolute growth rate of such a memoryless lineage *g*_*mless*_ = log (*N*_*mless*_(*t*_*max*_)*/N*_*mless*_(0)) from *g*_*m*_ to estimate the relative fitness of having memory *r* = (*g*_*m*_ *− g*_*mless*_)*/t*_*max*_.

We subject a growing population of cells to a set of 84 environmental sequences where bacteria are exposed to drug treatment sequences (each horizontal line in Fig. 3 A corresponds to a different environmental sequence). The total duration of the procedure is kept constant and the duration of drug treatment varies, with exposure to the permissive environment before and after drug exposure (also see SI). Using these environmental sequences, we compute for each condition the fitness of a set of lineages with different memory sizes. We observe a non-linear relationship between memory size and fitness (Fig. 3 B). While there is a general trend that more memory is beneficial when drug treatment increase, local maxima emerge at intermediate drug treatment lengths (Fig. SI2). This effect arises from the trade-off between producing a high proportion of ‘on’ cells (occurring with higher memories) and exiting rapidly from the ‘on’ state when the environment switches from drug to permissive (occurring with smaller memories). For extended treatment lengths, longer memory in such cases staves-off an inevitable population collapse.

However, memory is coupled to the transients as well as to the final frequency of ‘on’ cells observed at the end of each sequence. To disentangle the respective effect of these two factors, we compared the fitness of a lineage with memory (*n >* 1) to a lineage without memory (*n* = 1). To make lineages comparable, the memory lineage has a different leaching rate which results in the same equilibrium ‘on’ frequency under sustained treatment Fig. 3 C. The relative growth rate is the difference between the growth rate of a lineage with memory and without memory following http://myxo.css.msu.edu/ecoli/srvsrf.html. When the treatment length is short, longer memory is disadvantageous as compared to larger leaching rates. The disadvantage is because the cells stay in the ‘on’ state even after the short treatment has elapsed. For intermediate treatment lengths, more memory allows (i) to produce more ‘on’ cells and (ii) longer residual time in the ‘on’ state, then becoming advantageous. Yet, long residual times corresponding to longer lag times could conflict with the total sequence length where the fitness is measured. Hence, very long memory is also not useful as the cells then take much longer to exit the ‘on’ state as compared to the sequence length. Thus, we see the presence of local maxima in memories (driven partly by the absolute fitness of the memoryless lineage, see SI). Under lengthy sustained treatment, all lineages are at the ‘on’ equilibrium, and memory length is inconsequential. This analysis reveals that, for a range of conditions, memory outperforms ‘classical’ memoryless switching against an equal equilibrium value of ‘on’ cells.

## Discussion & Conclusion

Studies on phenotypic memory have typically focused on models assuming two distinct states, ‘on’ and ‘off’ [38, 12]. We show here that this is often used assumption hinders us from understanding the rich dynamics observed in experimental and empirical studies. We have addressed this gap between observation and theory by extending the analysis of the two type model to multiple underlying states.

Assuming two phenotypic observables (‘on’ and ‘off’) is already a simplification, however going forth with it, we have extended the underlying possible cellular states of the ‘on’ type. The number of ‘on’ states essentially acts as memory, since an increase in the number of compartments will delay the time required to get back to the ‘off’ state. Memory, defined as such, results in non-exponential distribution of residence time in the ‘on’ state [26].

Thinking of memory as a multi-state process appears to explain otherwise anomalous observations. At the single-cell level, the lag is the time it takes to resume division for a single cell taken out of a tolerant population that has just exited a prolonged antibiotic treatment [32]. Under artificial selection experiments, cells adapt their lag time to match treatment duration [19, 29] while displaying within-population variability. This variability increases with the mean lag time (see SI). Initially, this observation seems counterintuitive, as the best strategy would be for all cells to resume growth as soon as the treatment is over. [29] showed that heterogeneous lag times can promote survival to transient antibiotic treatments while having a negligible cost on population growth. From our model, we find that a cell in compartment *i* = *n*, …, 1 will experience a lag time (time to reach the off state where growth starts again) following a gamma distributed with shape parameter *i* and rate parameter *ϵ*. Incidentally, we note that the gamma distribution is a good model of lag times according to the experimental results [29].

Moreover, according to our framework, change the mean lag time of a cell population to match treatment length could be possible by a variety of ways: evolution in the memory length, in the leaching rate, in both or competition between lineages with variable properties of the gamma distribution. However, for a fuller understanding of the evolutionary forces acting on these traits, explicit considerations of their potential trade-offs, not included in our work, are required.

To some extent, the transients of the dynamics can be understood in terms of eigenvalue analysis. Complex eigenvalues introduce oscillations in the dynamics (SI). Larger memory sizes correspond to more solutions in the imaginary space which are reflected in the dynamics with more oscillations (Fig. 2). The magnitude of the effect hinges on how fast cells experience the memory (leaching rate) and the initial switch rate. For any leaching rate, however, as the memory increases, the oscillations (captured by the peak-to-peak amplitude) increase but only up to a limit (Fig. 2 B). At the population level, the difference is solely in the fraction of the ‘on’ cells and not in the population size. This result is based on our assumption of both the ‘on’ and ‘off’ cells having the same birth and death rates. Forgoing this assumption would lead to a further divergence between the composition and size of lineages with different memories. Whether fitness depends on composition or size of a lineage, memory will bring unique dynamical properties that might impact survival. Ecological context will set the timing of fitness evaluation, realising Darwinian selection on lineages (Fig. 2).

Phenotypic heterogeneity while advantageous for a lineage [25, 39] can be a nuisance when population expansion is harmful. Since Hobby and Bigger [36, 37] persisters have been a fly in the ointment for antibiotic treatment only exacerbated now by the antibiotic crisis. Similarly, in cancer, quiescent subclones are a persistent problem leading to relapse [40, 41, 42]. The subclone population dynamics also show over-shooting [43, 28]. We have presented the structure of the use of multi-state memory and its application to antibiotic tolerance. The persisters as defined in our case are a subpopulation of tolerant cells appropriately defined as “heterotolerant” [32]. While tolerance does not affect minimum inhibitory concentration, the duration of treatment will be crucial for the eradication of bacteria. Considering memory brings an additional time-scale which can be exploited for controlling pathogenic populations.

Numerous extensions of our approach are possible. For example, we have focused on heterogeneity resulting from only environmentally triggered switches, a property observed in several experimental models [44, 32, 3, 5]. Alternative switching mechanisms may depend on the intrinsic properties of the population such as density or composition. Similarly, the leaching mechanism is predetermined and constant across the compartments. It is possible that such processes are under complex, joint control of the organism as well as the environment. Although detailed experimental characterisation of bidirectional switching behaviors remains rare (due to technical challenges), we expect that memory-based switching is not an exception.

Oscillations involving excitation and decay can be identified across life. For example, excitable genetic switches are found in bacteria [45], and they belong to circadian clocks in cyanobacteria [46], in other non-photosynthetic bacteria [47], in Drosophila and in plants [49, 50]. The molecular mechanisms identified in these systems point towards a malleable duration of the oscillations by mutations [45, 51], modifications of the regulatory network [30] or degradation and destabilisation of protein complexes [48]. Given the explicit dependence on external signals, flies’ chronobiology has been proposed as a tool to understand adaptation to variable environmental conditions. Thus the adaptive evolution of molecular processes underpinning the oscillations of the excitable systems such as our model is conceivable.

While theoretical developments are essential, the applied aspect can be further exploited. Beyond antibiotics and cancer treatment, bioengineering and understanding of the formation of microbial consortia could be informed using our approach. Knowing the memory limitations (e.g. coming from decay rates of protein complexes) involved in critical oscillatory processes such as in chronobiology can inform us about the limits of adaptation under extreme environmental events or anthropogenic changes. Especially when harsh environments and time-lags are of importance such as in niche construction and the evolution of multicellularity [27, 52].

Understanding gene regulatory networks on a developmental landscape, a’ la Waddington [53], poses exponentially complex computational challenges (e.g. the explosion of possible attractors when considering multiple switches [54] and multiple phenotypes) (as also discussed in [55]). We show the existence of multiple local maxima for memory, that may change depending on the definition of fitness (population size or composition). This further forces us to rethink our concepts about possible phenotypic states and how they are determined by a plethora of molecular constructs (see SI for an extended discussion). How epigenetic memory functions over generations would focus on understanding how the phenotypic clock (molecules, appropriate histone modifications) are inherited. In our model, time as kept by degrading molecules is arbitrary. Inputs from chronobiology could help see how the molecular concentrations and circuits are entrained, resulting in biologically driven growth and leaching dynamics.

Phenotypic heterogeneity forces us to reassess the genotype-phenotype map in a fundamental manner. Choosing the appropriate phenotypic response to complex and varied environments is possible via a multitude of processes such as environmental sensing, epigenetic triggers and controlled molecular concentrations. Such processes that interpret the genetics to a large but finite phenotypic space to survive in a possibly infinite environmental space are extremely relevant for natural selection. Theories as described herein coupled with experiments exploring a number of environments will help us elucidate the variety of possible interpreting mechanisms bridging the genotype-phenotype divide.

## Acknowledgements

We thank Silvia De Monte and Orso Romano for helpful discussions. Funding from the Max Planck Society is gratefully acknowledged.

## Supplementary material

### Matrix model and solution

The system in Eq. 2 of the main text can be written in matrix form as

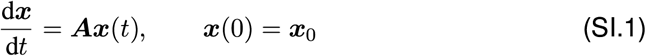

where

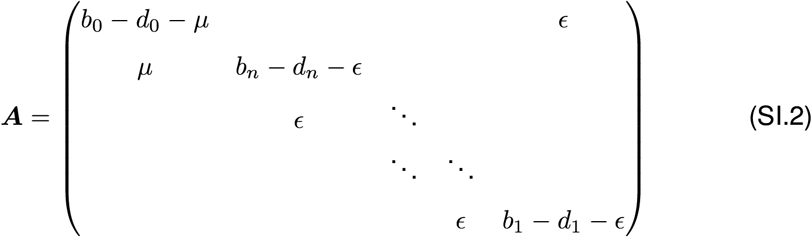

and ***x***(*t*) = (*x*_0_, *x*_*n*_, …, *x*_1_)^*T*^, here the superscript T indicates vector transposition. The solution to Eq. SI.1 is ***x***(*t*) = *e*^*t****A***^***x***_0_. The matrix ***A*** is essentially non-negative (i.e. off-diagonal entries are non-negative) and irreducible (i.e. any compartment can be reached from any other compartment). This ensures that ***A*** has a real dominant eigenvalue λ such that all other eigenvalues have smaller real part. As a consequence, asymptotically (*t →* ∞) the cell population grows exponentially with rate λ and the distribution of cells in the compartments approaches the right eigenvector ***u*** corresponding to λ, when this eigenvector is scaled so that its components add up to unity.

### Symmetric growth dynamics

Under complete symmetry in the growth dynamics for all compartments, i.e. *b*_*i*_ = *b* and *d*_*i*_ = *d*, we have that λ = *b − d*, i.e. growth equals birth rate minus death rate. Solving then the eigenvector equation ***Au*** = λ***u*** for the symmetric case, the equilibrium fraction of cells in any ‘on’ compartment (*i* = *n, n−*1, …, 1) is equal to *µ/*(*ϵ*+*nµ*).

**Figure SI.1:**
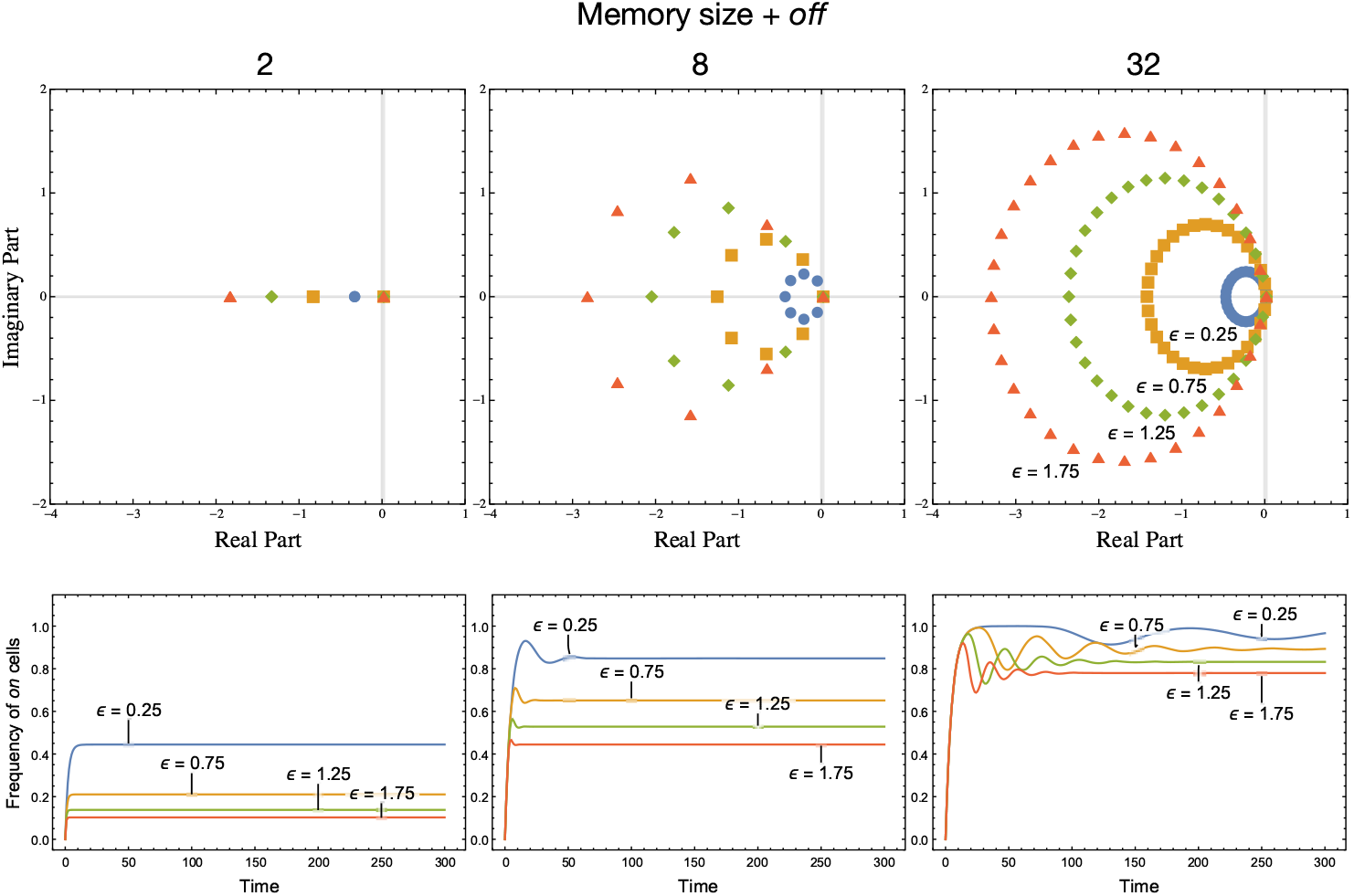
Eigenvalue spectra and population dynamics. (*b*_*i*_ = 1.0 and *d*_*i*_ = 0.98 for all *i* = 0, *n, n −* 1, …, 1) for different number of compartments (Memory *n* = 1, 7, 31 + 1) and different values of the leaching rate (*ϵ* = *{*0.25, 0.75, 1.25, 1.75*}*). The switching rate (*µ* = 0.1) is kept constant. This behaviour is related to the spectra of the matrix models (top) that govern dynamics (bottom), especially the overshooting.

Thus, the equilibrium fraction of cells in the ‘off’ state, i.e. in the zeroth compartment, is equal to *ϵ/*(*ϵ* + *nµ*). This expression shows that a change in the number of ‘on’ compartments and a change in the ratio of the switching rate to the leaching rate have the same effect on this equilibrium: if *n* or the *µ*-to-*ϵ* ratio increase (decrease), the equilibrium fraction of cells in the ‘on’ state increases (decreases). However, the equivalence breaks down when one looks at transient dynamics (Fig. SI.1). To understand why, let λ_0_, λ_1_, …, λ_*n*_ be the eigenvalues of ***A*** in Eq. SI.2 in order of decreasing real part and ***u***_0_, *u*_1_, …, ***u***_*n*_ be the corresponding eigenvectors, which are assumed linearly independent. Then, the solution to Eq. SI.1 can be written as

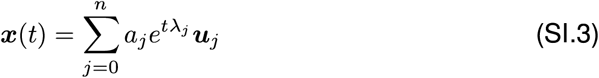

where *a*_*j*_ are fixed scalars corresponding to the coordinates of ***x***_0_ in the basis given by the eigenvectors of ***A***. This solution highlights how subdominant eigenvalues may influence dynamics for small *t*, i.e. before the asymptotic phase when these get dominated by λ_0_. The evolution of ***x***(*t*) relative to the influence of the dominant eigenvalue over time is given by 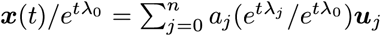. Since λ_0_ dominates the other eigenvalues, we have that 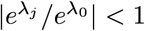 for *j >* 1 and we get that 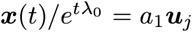 as *t →* ∞. But for small *t*, the quantities 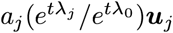 may be not negligible. Typically, the closer subdominant eigenvalues are in their real part to the dominant eigenvalue, the larger and more protracted the transients. Complex eigenvalues also introduce oscillations. In our case, as Fig. SI.1 shows, the more ‘on’ compartments are present, the more complex eigenvalues that are closer in real part to the dominant are found in the spectrum of ***A***. Dynamically, this produces greater overshooting and slower convergence. This is not observed when the *µ*-to-ϵ ratio increases yet a single compartment is present. For *n* = 1, the only subdominant eigenvalue is λ *− µ −* ϵ, which is real and gets closer to the dominant eigenvalue λ as the *µ*-to-ϵ ratio increases. But while this increase leads to an increase in the equilibrium fraction of ‘on’ cells, see formula above, no overshooting ensues, as Fig. 4 in the main text shows

### Environmental Variation

Populations of cells can respond to stressful environmental conditions in a variety of ways. We focus on sequences of environments composed of multiple seasons as a source of environmental stress. Each season is either conducive or harmful to the population. As a response, a lineage of cells is bestowed with different growth rates for the ‘off’ and ‘on’ compartments. The ‘off’ cells thus correspond to the growing, typically the observable quantity of a lineage. The ‘on’ cells do not grow but flow through the compartments. We look at ‘triggered’ persistence [32], i.e. the cells switch from ‘off’ to ‘on’ only when the season is harmful.

#### 5.1 Sequence diversity

As an example of controlled fluctuating environments, antibiotic treatment schedules are apt examples. Treatment regimes take various forms [19, 56]. A good understanding of the effect of sequence and even sequence memory of cells can help design effective treatment regimes aimed at minimising collateral resistance to multiple drugs.

In the main text we focused on a large number of treatment sequences (in total 84). The sequences ranged from no treatment to all treatment where each season lasted one time unit. The growth rates of lineages of cells with different memories was calculated as *g*_*m*_(*i*) = log(*N*_*tmax*_*/N*_0_)*/t*_*max*_ for each memory size *i*. We compare this Malthusian growth rate to that of a lineage which gives the same eventual ‘on’ frequency but for *n* = 1 i.e. a memoryless process (which will have a different leaching rate ϵ). Thus *r* = *g*_*m*_(*i*) *− g*_*mless*_(*n* = 1, ϵ)*/t*_*max*_ is the difference in the Malthusian growth rates of the two lineages if under direct competition (http://myxo.css.msu.edu/ecoli/srvsrf.html).

Using our minimal setup we can explore the effects of relaxing treatment for different lengths of time. However it would be useful to reduce the number of sequence to perform a thorough analysis. In the main text the fluctuating sequences last for 165 time-steps leading to 84 treatment regimes. We let each season last for 15 time-steps instead of 1, which yields us only 7 different sequences of 11 seasons each. The analysis of these sequences is shown in Fig. SI.3 and Fig. SI.4.

**Figure SI.2:**
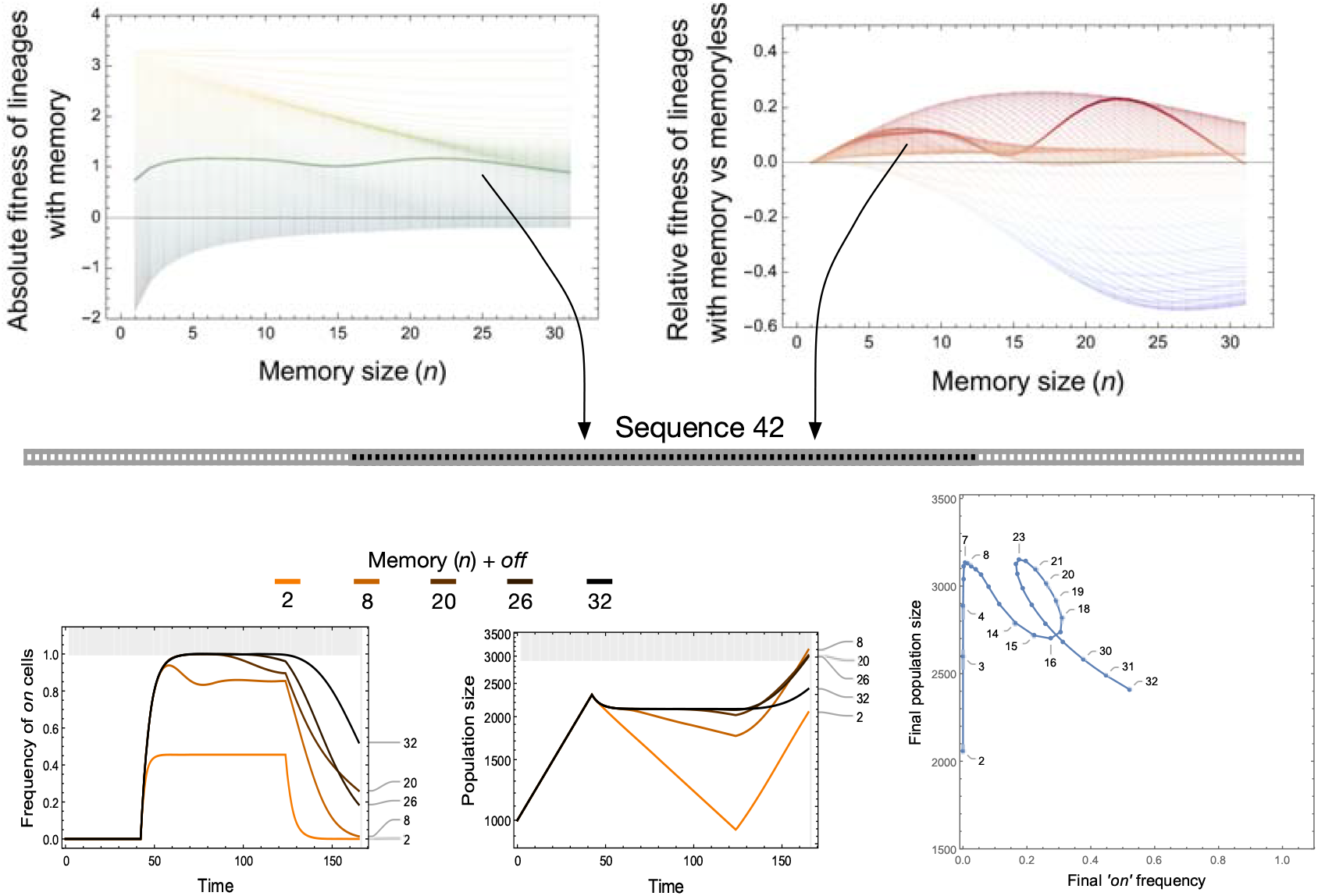
Eco-evolutionary details of a treatment regime. In the main text we explored 84 different sequences each lasting for 165 time-steps (Fig. 4). The sequences ranged from no treatment seasons to all treatment scenarios. The heat-maps in the main text, corresponding to the absolute and relative growth rate are explored in detail here in the top panels. For a specific sequence of seasons -sequence 42 -we show the the evolutionary factor, the frequency of ‘on’ cells and the ecological factor, the population size in the bottom panels. The results are shown for a chosen number of memory sizes (n) + 1 (‘off’ compartment). The non-linear relationship between the final ‘on’ cell frequency and the final population size is shown in the last panel as *n*+1 ranges from 2 to 32.

**Figure SI.3:**
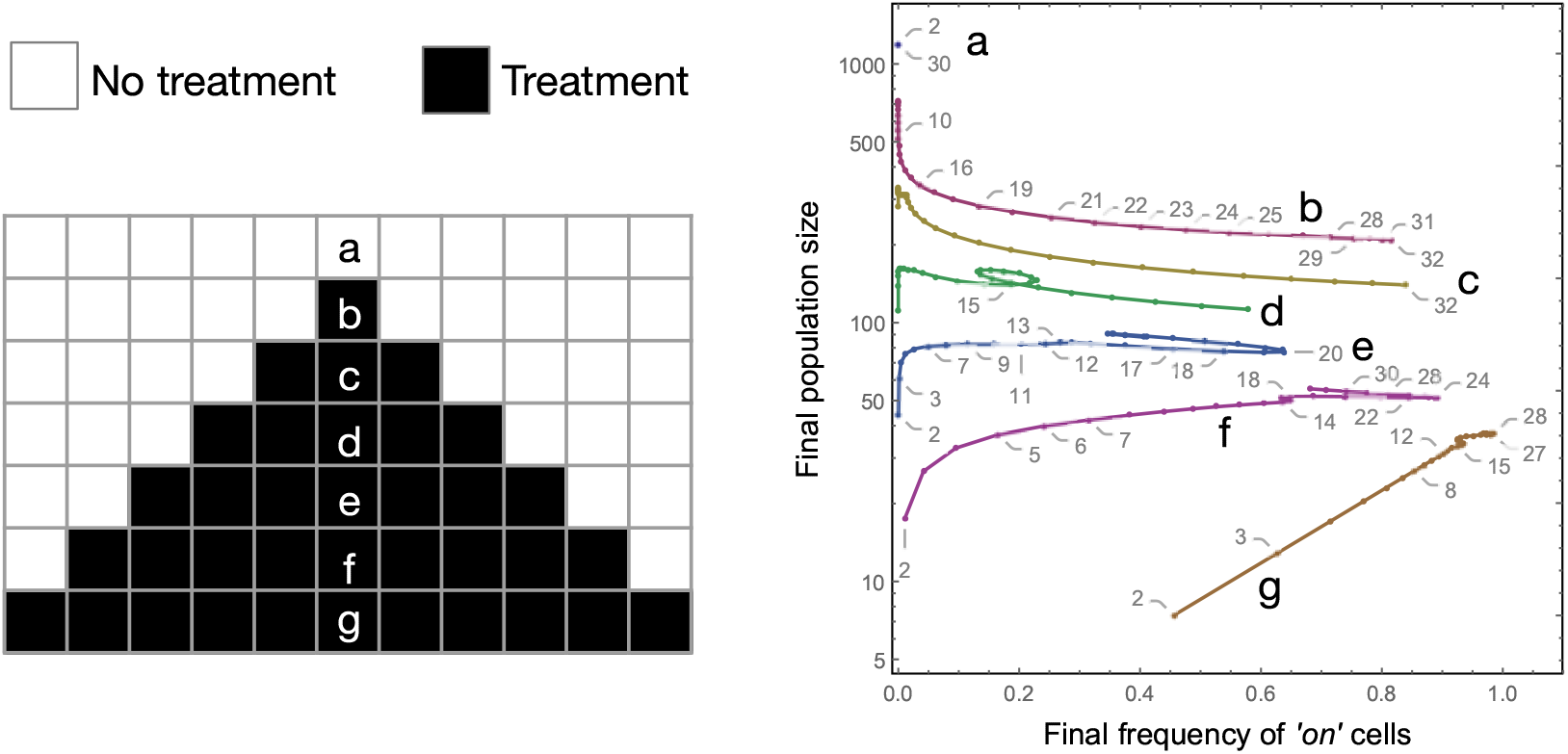
Final frequencies of the ‘on’ cells and the final population sizes of lineages with different memory lengths. The results of the population and evolutionary dynamics of the cell lineages with different memory sizes (*n* = (1, …, 31))+1 are plotted after different sequences of environments each comprising of 11 seasons. There are two kinds of seasons -treatment (filled) and no treatment (empty). Each season lasts for 15 numerical time-steps The sequences from (a) to (g) increase the length of antibiotic treatment where the ‘off’ cells die with rate *b*_*off*_ *− d*_*off*_ = 1 *−* 1.02 = *−*0.02. The ‘on’ cells do not grow. In the absence of antibiotics there is no switching from ‘off’ to ‘on’ i.e. *µ*_no treatment_ = 0 whereas treatment triggers the switch *µ*_treatment_ = 0.2.

#### 5.2 Condition dependent switching

In the main text we have assumed ‘triggered persistence’ i.e. when not under treatment the switch from ‘off’ to ‘on’ does not work. Relaxing this assumption, in Fig. SI.5, we show the result of the dynamics on the spectrum of constitutive switching where (*µ*_treatment_ = *µ*_no treatment_ = 0.2) to strictly antibiotic triggered switching (*µ*_treatment_ = 0.2, *µ*_no treatment_ = 0.0). If a lineage switches to the ‘on’ cells constitutively then the eventual population size is much smaller as the lineage loses cells to the non-growing state and the growth can only proceed after they have exited the ‘on’ cells after leaching through all the memory states. The exact shape of the plotted clines depend further on the specific sequence chosen to be explored.

**Figure SI.4:**
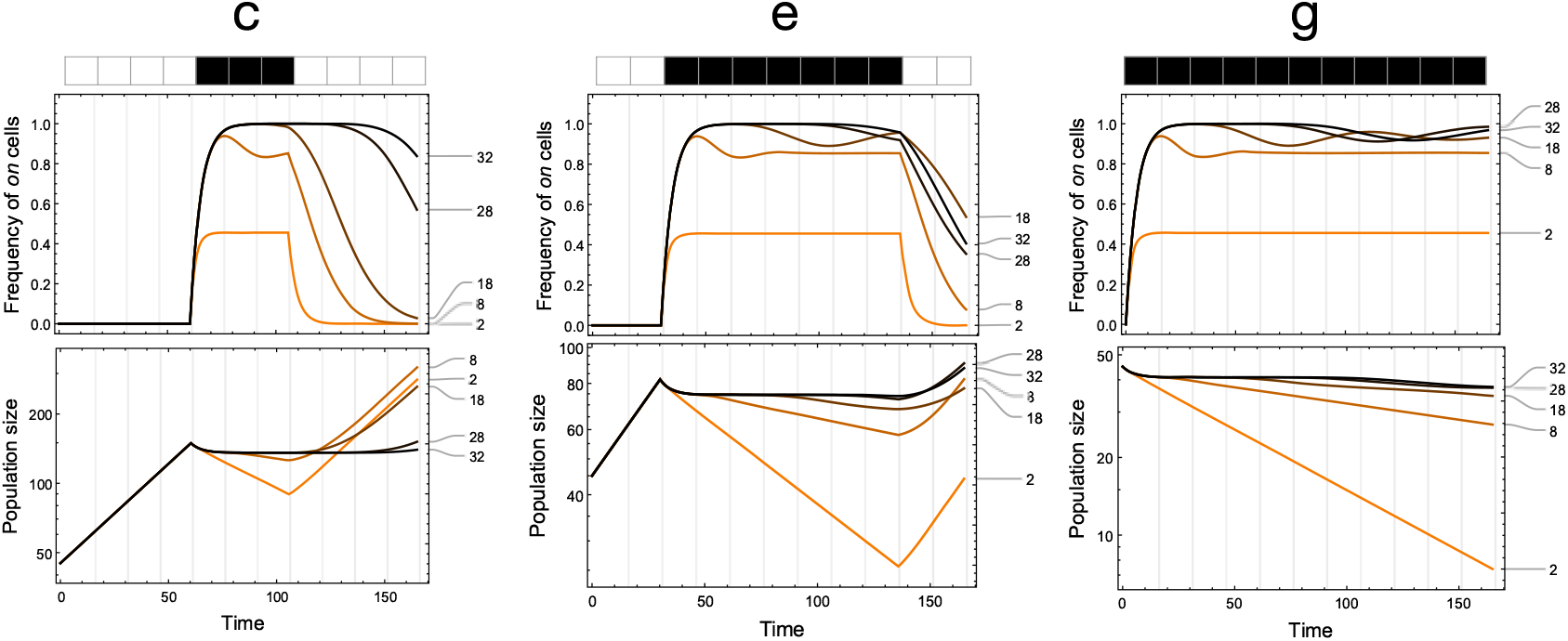
Eco-evolutionary dynamics of lineages with different memories under select treatment regimes. To better understand the non-linear behaviour seen in the Fig. SI.3 we choose three different treatment regimes (c), (e) and (g). The plot denote the temporal dynamics of (top) the fraction of ‘on’ cells and the (bottom) population size for lineages with different memory sizes (*n* + off = 2, 8, 18, 28, 32). Under no treatment (empty) the ‘off’ cells grow with rate *b*_*off*_ *− d*_*off*_ = 1 *−* 0.98 = 0.02 and there is no switching from ‘off’ to ‘on’ (*µ*_no treatment_ = 0). Under antibiotic treatment the ‘off’ cells die with rate *b*_*off*_ *− d*_*off*_ = 1 *−* 1.02 = *−*0.02 and switching occurs with rate *µ*_treatment_ = 0.2. The ‘on’ cells do not grow. For treatment sequence (c) we have a short treatment length. The memoryless lineage (*n* + 1 = 2) reaches equilibrium in ‘on’ cells immediately. Lineage with 8 compartments shows a characteristic over and undershoot in the short treatment time whereas the others have longer amplitudes and the treatment is not long enough to see them. When the treatment is finished the shorter memory lineages have a shorter time lag in resuming growth. For treatment sequence (e) the transients of the longer memories show the over and undershoots affecting the final on proportions. Also since there in not much time left after the treatment ends, the shorter memory lineages do not have enough time to increase in population size. Under sustained treatment in (g) the ‘on’ cell equilibrium is being approached but the sequence length is not enough for all lineages to achieve it. Only the memoryless and the lineage with 8 compartments reach equilibrium while the others still show oscillations. Under sustained treatment the lineages decline in population size but a larger memory buffers the time of decline. The initial population consist of all 45 all ‘off’ cells. Each season (treated or untreated) lasts for 15 numerical time-steps. The leaching rate is set to ϵ = 0.25.

#### 5.3 Long term growth dynamics for cyclic environments

Using the analysis in [13], we also consider long term dynamics when the population goes through the same seasonal sequence a repeated number of times. We form two matrices ***A***_R_ and ***A***_T_ with the same structure as in Eq. SI.2. Matrix ***A***_R_ is parametrized to reflect the growth and phenotypic dynamics of the population under a relaxed season, while matrix ***A***_T_ is parametrized to reflect the growth and phenotypic dynamics of the population under a treatment season. Given an initial population composition ***x***(0), the final population composition ***x***(*nτ*) after a sequence comprising *n* seasons each lasting *τ* units of time is obtained as

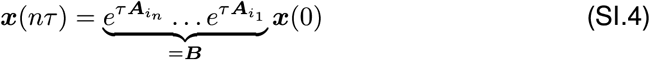

where *i*_*j*_ ∈ *{*R, T*}, j* = 1, …, *n*. When the population cycles *m* times through the same sequence, the population composition at the end of the *m*-th cycle is ***x***(*nτm*) = ***B***^*m*^***x***(0). As *m* gets large, the population size grows by a factor equal to the dominant eigenvalue of ***B*** after having gone through a sequence. Using this approach, it can be shown (Fig. SI.6) that short term population growth for different memory sizes as explored in Fig. 5 of the main text displays a strikingly qualitatively similar behavior as long term growth.

However, ***B***, as a product of matrices, has eigenvalues that are invariant to how matrices are arranged in this product. Therefore, this approach does not extend to our analysis in the next section where the effects of season permutations are explored.

**Figure SI.5:**
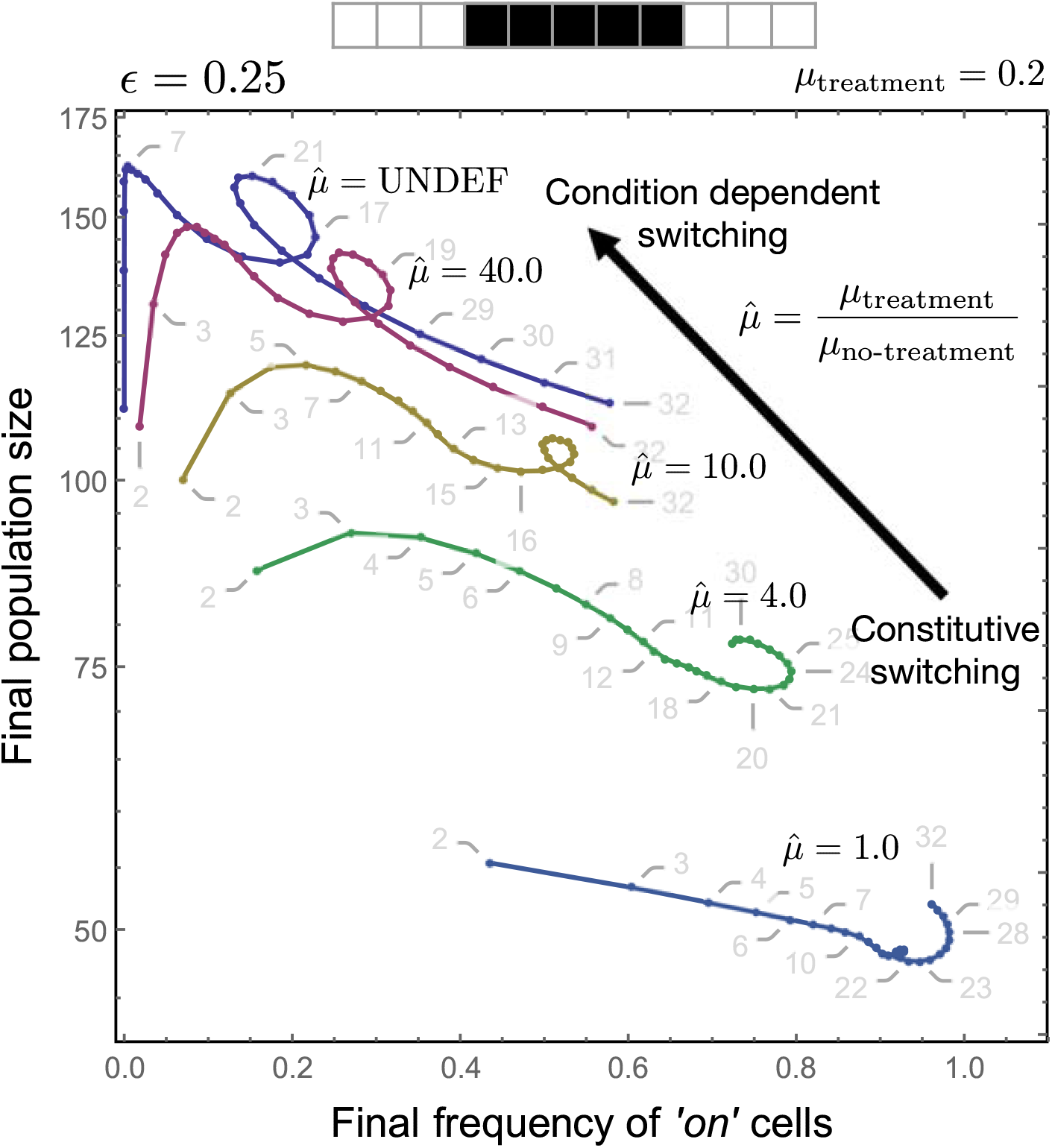
Constitutive switching to “triggered” persistence. For the antibiotic tolerance we assumed that the ‘on’ cells are produced only when under treatment (*µ*_*treatment*_ = 0.2 and *µ*_*no−treatment*_ = 0.0). Here we explore the effect of relaxing this assumption. We define 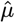 as the ratio between *µ*_*treatment*_ and *µ*_*no−treatment*_. For constitutive switching i.e. switching irrespective of the environment, the ‘on’ cells can saturate for longer memories, however since the ‘on’ cells do not grow, the population sizes are low. As we move towards condition dependent switching, the population is rescued with the final population size showing a prominent non-linear dependence on the memory size *n* (grayed out numbers showing *n* + 1). Also since ‘on’ cells are produced only under treatment and they do not grow, the final frequency of ‘on’ cells is lower than when under constitutive switching.

**Figure SI.6:**
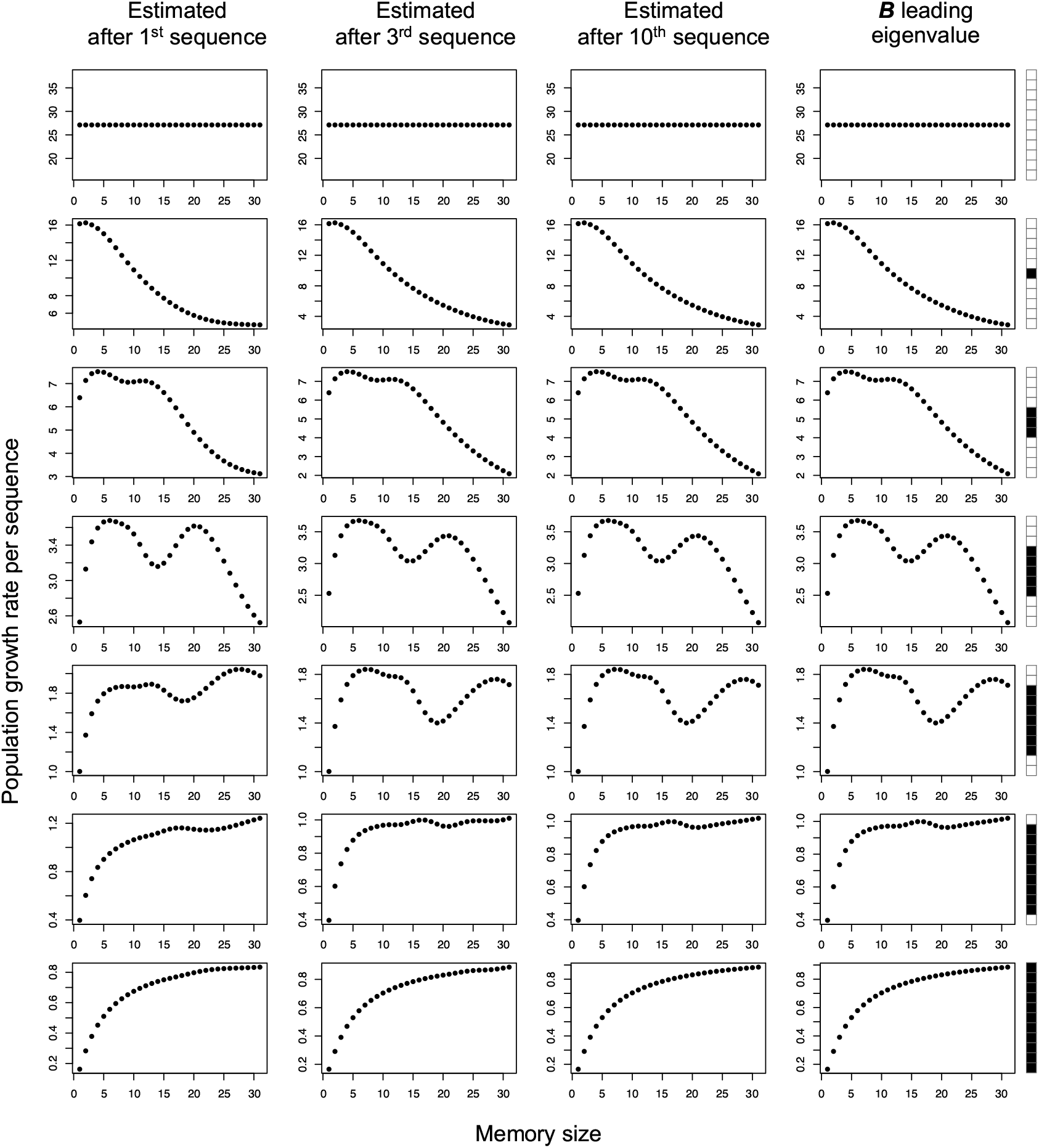
Long term growth dynamics for cyclic environments. Using the same parameters as in Fig. SI.3, growth rate (y-axis) is estimated as the factor by which population has grown after having gone through one sequence of seasons. Cycling through the same sequence, the estimation is done after the first cycle, the third and the tenth. Resulting growth rates are compared with those expected in the long run as given by the leading eigenvalue of the matrix ***B*** that captures population dynamics over a sequence.

#### 5.4 Lag time distributions

Upon sufficiently long exposure to sustained treatment, the population reaches a stable compartmental distribution given by the right dominant eigenvector ***w***_T_ = (*w*_0_, …, *w*_*n*_)^*T*^ of the ***A***_T_ matrix scaled so that the eigenvector components add up to 1. This enables us to compute the lag time distribution when cells in ‘on’ state have no growth dynamics, i.e. *b*_*i*_ *− d*_*i*_ = 0 *−* 0 for *i* = 0, …, *n*, as assumed above. A randomly sampled cell from the exposed population is in compartment *i* with probability *w*_0_. When the cell is in the zeroth compartment, i.e. ‘off’ state, its lag time is 0, as it is already dividing. When the cell is in any ‘on’ compartment, i.e. *i* = 1, …, *n*, its lag time is the time it takes for the cell to reach the ‘off’ state, i.e. the zeroth compartment, where division occurs. This time *τ* is gamma distributed with shape parameter *i* and rate parameter ϵ so that, a cell from compartment *i* has lag time *τ* with probability Gamma(*i*, ϵ, *τ*). The probability that a sampled cell from the treated populations has lag time *τ* is then given by,

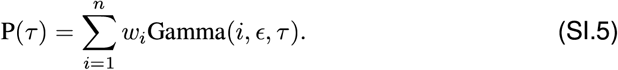

#### 5.5 Permutations

Another way of designing treatment regimes is to permute a given number of treatment seasons. As a template we look at the permutations of 4 treatment seasons and 4 relaxed seasons for a total of 70 possible sequences. We check the final size of a lineage once it has experienced all 8 seasons. We do so for cell lineages with different memory lengths. Plotting the final population size against the final ‘on’ cells frequency we see that an increase in the number of compartments typically results in an increase in the final ‘on’ frequency (but not always, see the loops), Fig. SI.7. Also the population size peaks typically at intermediate memory length. An intermediate memory size is then advantageous when selection operates on the surviving population at the end of the treatment sequence.

**Figure SI.7:**
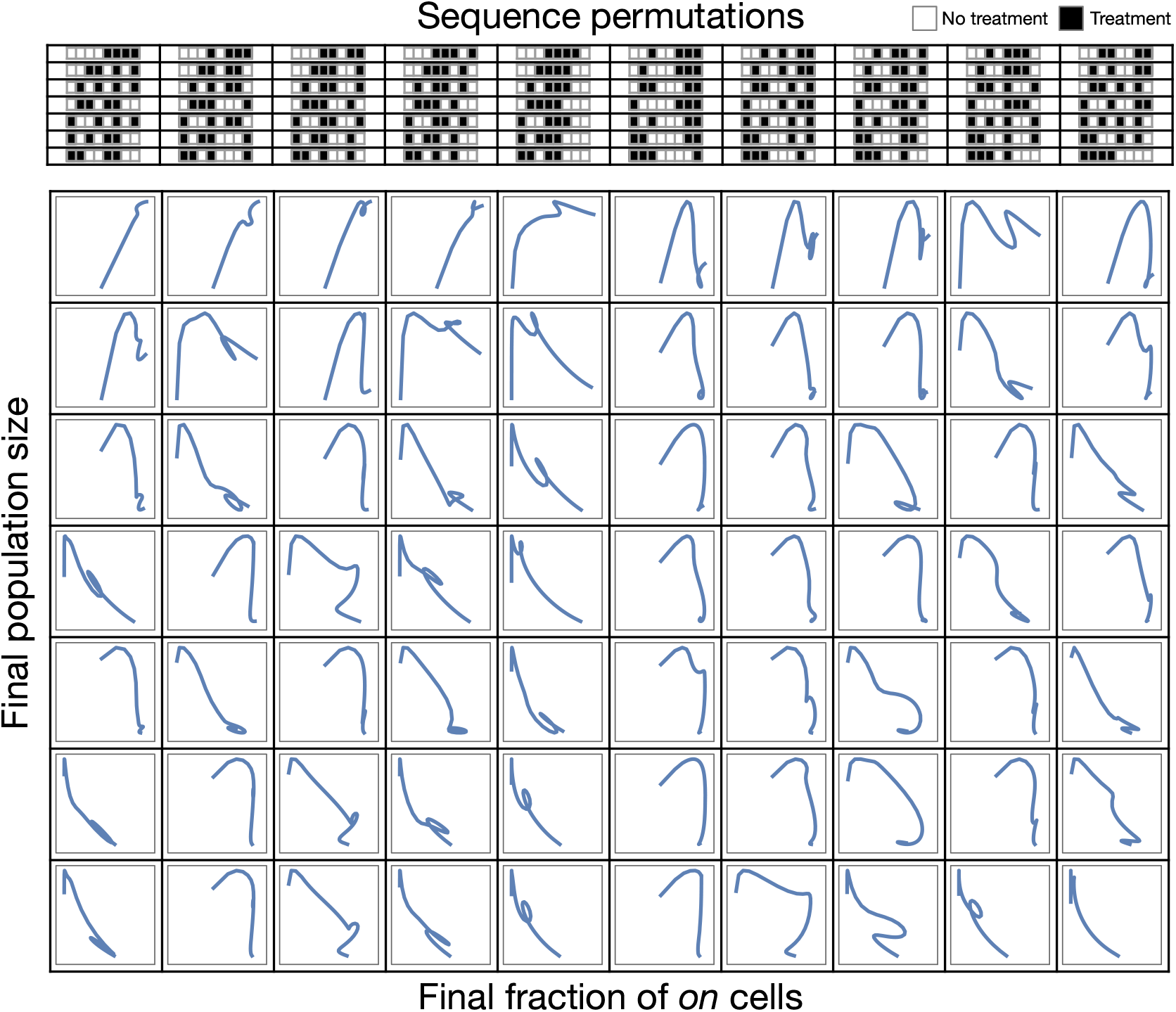
Eco-evolutionary outcomes for sequence permutations. For the 70 different permutations of the sequences consisting of 4 treatment seasons and 4 no treatment (so permutations of 1,1,1,1,2,2,2,2). Under condition dependent switching (*µ*_treatment_ = 0.2 else 0), parameters are (*n* = (1, …, 31)) *b*_*off*_ *− d*_*off*_ = 1 *−* 0.98 = 0.02 under no treatment and *b*_*off*_ *− d*_*off*_ = 1 *−* 1.02 = *−*0.02 under treatment. Each season (treated or untreated) lasts for 15 numerical time-steps. Leaching is constant at rate ϵ = 0.25. Overall, we see that numerous sequences peak in final population for intermediate memory sizes and non-linearity is a general observable.

### Phenotypic determinism

Phenotypes can be determined jointly by the internal states of a cell (e.g. intracellular protein concentrations) and external effects (available metabolites). Furthermore, the number of possible internal determinants can be numerous. Irrespective of the the possible complexity of phenotypic determinism, in our study we have assumed a system which is best depicted by Fig. SI.8 (a). A trigger (ecological-biotic or abiotic) or a stochastic process is assumed to increase the concentration of a certain phenotypic determinant that can be tracked. We have assumed only two phenotypic states -‘on’ and ‘off’. Also the decay landscape is gradual. As shown in Fig. SI.8 (a) it is possible that the decay process of the determinant is not smooth and (b) can lead to massive variation in the time spent in the two states. Fig. SI.8 (c) highlights the case when the concentration can determine the same two phenotypes in a redundant fashion. Both very low and very high concentrations generate the same phenotype ‘off’ whereas the intermediate is the ‘on’ state. Indeed an exact distinction between phenotypes is limited by the tests possible to differentiate between the phenotypes and their precision. Multiple phenotypes as shown in Fig. SI.8 (d) when considered will disrupt our classical analysis but will need to be included as molecular tools get more and more precise.

**Figure S8:**
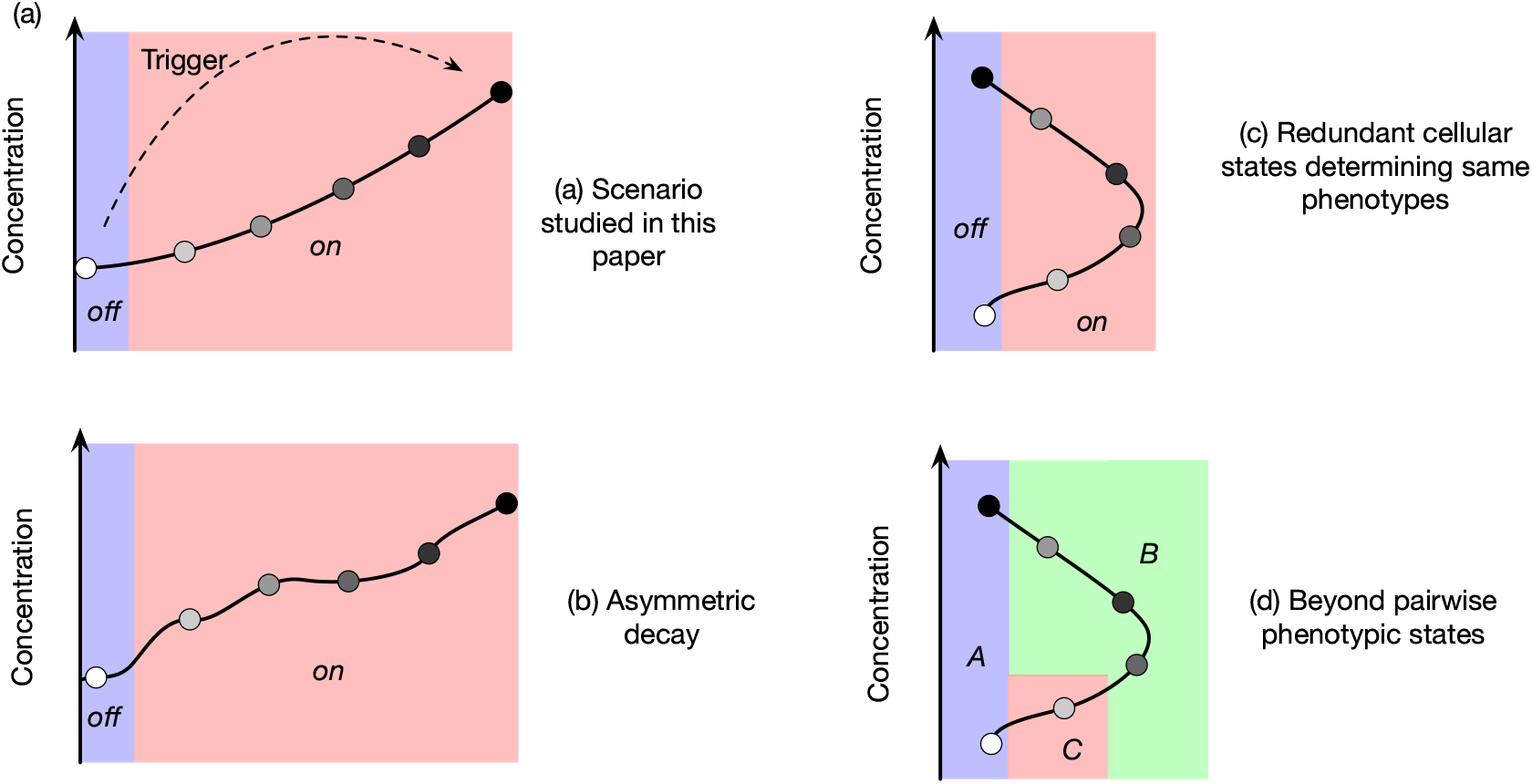
Some complexities in phenotypic determinism. An intracellular product, the presence and concentration of which depend on contingent ecological conditions, is a determinant of phenotypic state. (a) In our study we have assumed the simplest case where there are only two phenotypes, one of them characterised by multiple states. The decay of the determinant is smooth between the observed states and hence leads to tractable analytics. (b) It can be possible that the decay between the different tracked states is not smooth and this will affect the eventual distribution of phenotypes and the time spent in the two states. This situation can easily be captured in our framework by setting state dependent leaching rates, i.e. ϵ _*i*_, in Eq. SI.2. (c) Even if the decay is smooth it is quite possible that the determinism incorporates redundancy. In this example, both low and high concentration leads to the same phenotype. To include this scenario in our framework, we envisage the addition of an additional state *n* + 1 that also leads to the ‘off’ phenotype and from which cells can switch to the previous state *n* acquiring the ‘on’ phenotype. (d) While two phenotypes are easier to handle both experimentally and theoretically, this is probably the result of studying two environments. Multiple phenotypes are a reality and for a complete understanding of phenotypic heterogeneity, multiple phenotypes need to be incorporated, as binary cases are often special cases.

